# Metazoan-like kinetochore arrangement masked by the interphase RabI configuration

**DOI:** 10.1101/2020.09.09.289066

**Authors:** Alberto Jiménez-Martín, Alberto Pineda-Santaella, Daniel León-Periñán, David Delgado-Gestoso, Laura Marín-Toral, Alfonso Fernández-Álvarez

## Abstract

During cell cycle progression in metazoan, the kinetochore, the protein complex attached to centromeres which directly interacts with the spindle microtubules, the vehicle of chromosome segregation, is assembled at mitotic onset and disassembled during mitotic exit. This program is assumed to be absent in budding and fission yeast because kinetochore proteins are stably maintained at the centromeres throughout the entire cell cycle. In this work, we show that the assembly program at the mitotic onset of the Ndc80 complex, a crucial part of the outer kinetochore, is unexpectedly conserved in *Schizosaccharomyces pombe*. We have identified this behavior by removing the Rabl chromosome configuration during interphase, in which centromeres are permanently associated with the nuclear envelope beneath the spindle pole body. Hence, the Rabl configuration masks the presence of a program to recruit Ndc80 at mitotic onset in fission yeast, similar to that taking place in metazoan. Besides the evolutionary implications of our observations, we think that our work will help understand the molecular processes behind the kinetochore assembly program during mitotic entry using fission yeast as the model organism.

## Introduction

The three-dimensional configuration of the genome in yeast is characterized by the evolutionarily conserved Rabl chromosome configuration, defined by the stable association of centromeres and telomeres to the nuclear envelope (NE) (Jin et al. 1998; Jin, Fuchs, and Loidl 2000; Taddei and Gasser 2012). The NE comprises the inner nuclear membrane (INM) and the outer nuclear membrane (ONM), wherein the INM proteins play key roles in the interaction of the NE with chromatin (Czapiewski, Robson, and Schirmer 2016; Fernandez-Alvarez and Cooper 2017a). In particular, in fission yeast, centromeres are clustered together at the INM beneath the spindle pole body (SPB, the centrosome equivalent in yeast) and opposite to the nucleolus (Funabiki et al. 1993; Jin et al. 1998; Jin, Fuchs, and Loidl 2000), in a kinetochore-dependent manner. The linkage between centromeres and the INM occurs via the SPB-associated LINC complex (the <u>l</u>inker of <u>n</u>ucleoskeleton and <u>c</u>ytoskeleton), which comprises the KASH-domain ONM proteins (Kms1 and Kms2) and the SUN-domain INM protein Sad1(Hagan and Yanagida 1995; Hiraoka and Dernburg 2009; Unruh, Slaughter, and Jaspersen 2018; Shimanuki et al. 1997), which plays an essential role in supporting the kinetochore-SPB association. Besides, this connection is strengthened by the protein Csi1, which bridges Sad1 and outer kinetochore proteins, together with the conserved HEH/LEM domain INM protein, Lem2, which localizes at the nuclear periphery and the SPB (Hou et al. 2012; Barrales et al. 2016; Fernandez-Alvarez and Cooper 2017a). It is thought that the Rabl configuration reflects the positioning of the chromosomes during their segregation from the preceding mitosis which, in metazoan, is dismantled at the mitotic exit, but, in yeast, it is maintained throughout the subsequent interphase (Therizols et al. 2010). The reasons why the Rabl chromosome configuration in yeast is not disassembled at the mitotic exit and is maintained throughout interphase have been a mystery during decades. Recently, it has been observed in fission yeast that the interaction of at least one centromere with the SPB during interphase is a prerequisite to trigger the SPB insertion into the NE, a crucial event to nucleate the spindle microtubules. Hence, the destruction of the Rabl chromosomal organization abolishes SPB insertion and spindle formation and, consequently, leads to cellular lethality (Fernandez-Alvarez et al. 2016).

Attachment of centromeres to the SPB and, thus, the maintenance of the Rabl configuration, is supported by the kinetochore, the protein complex built on them (Cheeseman 2014). This large protein complex comprises around 80 members identified in humans, and its major components are conserved throughout eukaryotes. It can be subdivided into two distinct regions: the inner kinetochore, which forms the interphase with chromatin, and the outer kinetochore that constitutes the platform for interacting with spindle microtubules. Therefore, the kinetochore establishes the chromosomal attachment place for spindle microtubules, the motor that drives chromosomes distribution into daughter cells (Cheeseman and Desai 2008).

Importantly, kinetochore composition is dynamically regulated during the cell cycle in metazoan (Hara and Fukagawa 2018). There are kinetochore proteins that are constitutively present at centromeres (centromere-associated network, CCAN), but most of them are recruited to the kinetochore during late G2, prophase, or mitosis; this manner, members of the outer kinetochore complexes, for instance, Mis12 and Ndc80, are precisely recruited to the centromeres in late interphase and during prophase, respectively; consistently, once chromosomes are segregated, Ndc80 and Mis12 are orderly depleted following the onset of anaphase or the end of mitosis, respectively (Cheeseman and Desai 2008; Dhatchinamoorthy, Mattingly, and Gerton 2018; Nagpal and Fukagawa 2016). This well-regulated kinetochore components recruitment to the centromeres is assumed to be absent in *S. pombe*, where most of the outer kinetochore components such as Ndc80 and Nuf2, are constitutively present at centromeric regions throughout the cell cycle (Biggins 2013; Wigge and Kilmartin 2001) and only the components of the DASH complex, an essential element of the kinetochore that is required for the bi-orientation of sister chromatids, are recruited during mitosis (Liu et al. 2005; Cheeseman et al. 2001; Janke et al. 2002).

Two hypotheses try to explain the absence of the outer kinetochore recruitment program at mitotic onset in fission yeast: firstly, the absence of the nuclear envelope breakdown (NEBD) (Gu, Yam, and Oliferenko 2012); it has been observed that outer kinetochore disassembly/assembly program is a mechanism coordinated with the NEBD in metazoan (Hattersley et al. 2016); thus, the fact that NE is not disassembled before mitosis in *S. pombe* suggests that it might not be an efficient mechanism of outer kinetochore formation which it would involve an active transit of proteins from the cytoplasm to the nucleus. Secondly, the preservation of the outer kinetochore structure during interphase might be justified by its crucial role of maintaining the Rabl configuration (Asakawa et al. 2005; Takahashi, Chen, and Yanagida 2000); thus, in this case, the absence of an assembly program in fission yeast mitosis would be, in principle, independent of the absence of a proper NEBD.

It is challenging to discern whether the absence of the NEBD or the outer kinetochore role in the Rabl configuration is the reason behind the absence of the outer kinetochore disassembly/assembly cycle in fission yeast. This problem would be enlightened by studying the behavior of the kinetochore in cells without the interphase Rabl chromosome configuration. However, the idea of generating Rabl-deficient cells without compromising neither kinetochore structure nor cell viability has been challenging to address during the last decades. For instance, mutations on Nuf2 or Ndc80 partially remove Rabl configuration but also alter kinetochore structure (Asakawa et al. 2005; Hsu and Toda 2011; Nabetani et al. 2001); on the other hand, the presence of a thermosensitive allele of *sad1* (*sad1.2*) at restrictive growth temperature (36°C) abolishes all centromere-SPB associations but immediately leads to cell death (Fernandez-Alvarez et al. 2016). Hence, the development of a new system where the intact kinetochore was completely disconnected from the SPB at 32°C (standard temperature of growing for fission yeast) without dramatically impairing cell viability would help further disclose the behavior of kinetochore proteins in the absence of the Rabl nuclear configuration during yeast interphase. With the stated purpose, we here identified that the combination of the *sad1.2* allele together with the deletion of *csi1* at the semi-permissive temperature of 32°C generates severe centromere dissociation defects. In contrast, most of the cells are still viable due to occasional centromere interaction with the SPB, which is sufficient to trigger spindle formation. This manner, *sad1.2 csi1*Δ is a new scenario in which it is possible to characterize the behavior of the kinetochores dissociated from the SPB independently of its essential function of maintaining the Rabl chromosome configuration.

In this paper, we found that the dissociation of the centromeres from the SPB induces outer kinetochore disassembly during interphase and, more unexpectedly, we showed that, similar to metazoan, the outer kinetochore is reassembled in late G2. These results suggest that the outer kinetochore assembly program at mitotic onset coordinated with cell cycle progression is conserved in fission yeast, so far not observed because of being masked by the Rabl configuration. Our observations place *5. pombe* as a model organism to study the mechanisms behind the kinetochore assembly program highly conserved in metazoan and with enormous relevance for faithful chromosome segregation during cell cycle progression.

## Results and Discussion

### Levels of Ndc80 and Nuf2 at the centromeres remain constant throughout mitotic interphase

To accurately address the kinetochore proteins behavior at centromeres during the cell cycle progression in fission yeast, we first followed the focal intensity of endogenously GFP-tagged proteins on exponentially growing wild-type cells by live fluorescence microscopy, which also allowed to discard possible signal fluctuations. We analyzed Mis6 and Cnp20 (CENP-T ortholog) as representative members of the inner kinetochore (Hou et al. 2012; Takahashi, Chen, and Yanagida 2000) and Mis12, another outer component of the NMS complex (Obuse et al. 2004), which is also assembled and disassembled at mitotic onset and exit, respectively, during the metazoan cell cycle, but constantly attached to centromeres in yeast (Biggins 2013). Besides, we also studied the levels of Ndc80-GFP and Nuf2-GFP (Ndc80 complex) as part of the outer kinetochore, that localize at centromeres during interphase in budding and fission yeast (Biggins 2013; Liu et al. 2005). All images were processed at these experiments, and the protein intensity levels were quantified (see Methods section). Using this approach, we delineated the focal intensity of all these kinetochore proteins normalized per SPB signal (visualized via Sid4-mCherry) (Figure 1). As expected, our results indicated that all kinetochores components appear in a conspicuous focus in interphase at the SPB due to the stable centromere-SPB associations. Spindle formation (visualized by ectopically expressed mCherry-Atb2, tubulin) occurred for all analyzed kinetochore proteins during mitosis (Figure 1). Moreover, the focus intensity of all studied kinetochore proteins showed stable presence without significant signal fluctuation throughout interphase (Figure 1); in particular, Ndc80 and Nuf2, the two outer kinetochore canonical proteins, which modified their recruitment throughout the metazoan cell cycle, stably colocalized with the SPB signal during the experiment (Figure 1D and 1E). Our results discarded significant microvariations in all analyzed kinetochore protein signals, showing their recruitment to centromeres and maintaining their interactions with SPB along the whole cell cycle, which confirm the existence of a differential behavior between the fission yeast and metazoans outer kinetochore proteins. The prominent role of Ndc80 and Nuf2 supporting the Rabl chromosome configuration might be behind this fact. Thus, to explore this hypothesis, our efforts focused on removing the Rabl nuclear organization, neither compromising cell viability nor altering the stability of the kinetochore complex to evaluate the recruitment program of kinetochore proteins in the absence of their SPB associations at normal growth conditions.

**Figure 1.**
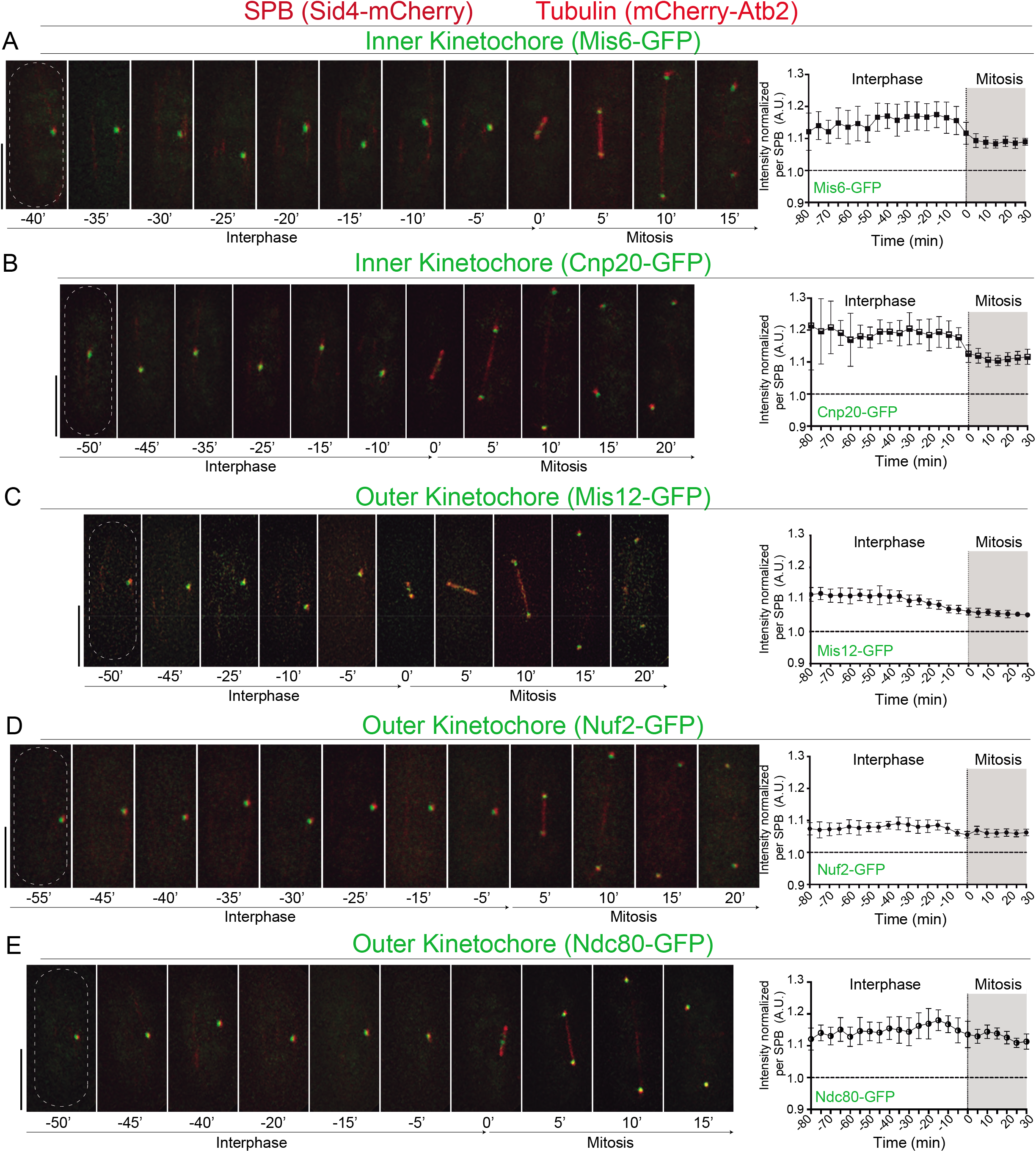
Outer kinetochore components Ndc80-GFP and Nuf2-GFP signal intensities at the SPB stay stable throughout interphase. (A-E) (Left panels) Frames from films of cells carrying Sid4-mCherry (endogenously tagged; SPB), ectopically expressed mCherry-Atb2 (controlled by *nda3* promoter; Tubulin) and GFP-endogenously tagged Mis6, Cnp20, Mis12, Nuf2 and Ndc80 as different kinetochore markers. Numbering indicates mitotic progression in minutes; t = 0 means first frame after SPB duplication. Bars, 5 μm. (Right panels) Mean Mis6-GFP, Cnp20-GFP, Mis12-GFP, Nuf2-GFP, and Ndc80-GFP intensities at the SPB throughout interphase and mitosis were quantified for each kinetochore marker (N=10 each). Error bars represent standard deviation. The data shown are from more than three independent experiments.

### Neither nuclear microtubules nor actin plays an essential role in maintaining the Rabl configuration in fission yeast

We explored different approaches to remove the interphase kinetochore-SPB associations at 32°C without altering the stability of both protein complexes. In budding yeast, centromere association to the NE requires nuclear microtubules (Bystricky et al. 2004; Jin, Fuchs, and Loidl 2000). For this reason, we first explored the possible role of microtubules in maintaining the Rabl configuration in fission yeast. To study the role of microtubules on centromere association to the SPB, we added the microtubule-depolymerizing drug carbendazim (MBC) to exponentially growing *wt* cells harboring Sid4-mCherry and mCherry-Atb2 as SPB and tubulin markers, respectively. Centromeres were visualized by endogenously GFP-tagged of the inner kinetochore protein Mis6. These experiments showed that the addition of MBC at concentrations able to completely eliminate microtubule formation did not induce any obvious centromere clustering defects (Figure 2A). To strengthen this observation, we quantified the distance between SPB and centromeres in growing cells with and without the addition of MBC, measuring the distance between Sid4-mCherry and Mis6-GFP foci (see Methods section) (Figure 2C–2E). Using this approach, we did not find any significant difference between both conditions, suggesting that, in contrast to *S. cerevisiae*, the maintenance of the Rabl chromosome configuration in *S. pombe* might not depend on nuclear microtubules polymerization (Jin, Fuchs, and Loidl 2000).

**Figure 2.**
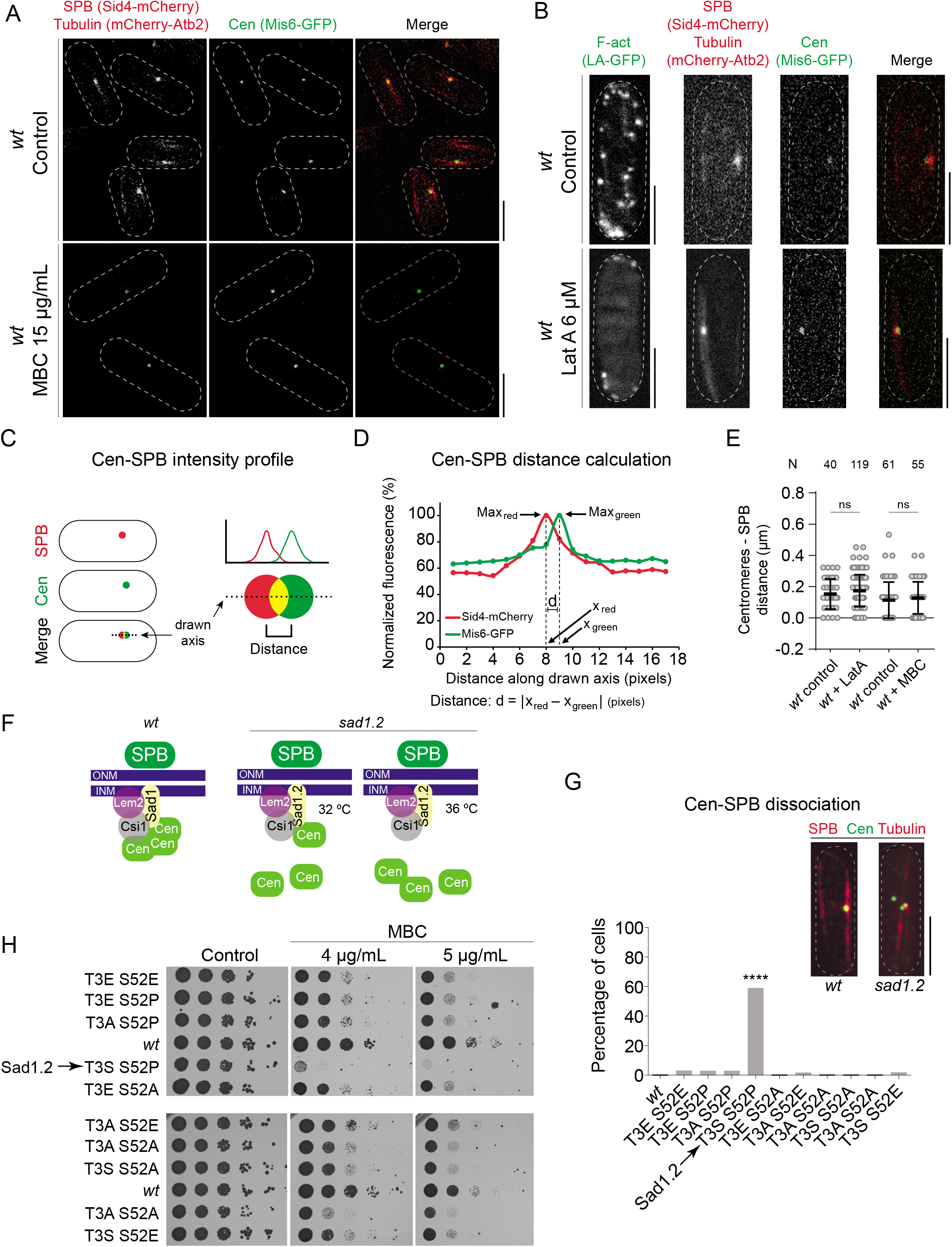
The interphase Rabl chromosome configuration in fission yeast is independent of microtubules and actin. (A-B) Live fluorescent microscopy images of *wt* interphase cells harboring Mis6-GFP and SPB and tubulin tagged as in Figure 1. (A) Top panel: in interphase, centromeres stay clustered together colocalizing with the SPB. Bottom panel: addition of the microtubule-depolymerizing drug carbendazim (MBC, 15 μg/mL) leads to loss of microtubules signal (proving microtubules depolymerization and thus efficacy of the MBC treatment), but the clustering of centromeres and co-localization with the SPB remains identical to that seen in a control setting. (B) A similar treatment was performed using Latrunculin A (Lat A, 6 μM). F-Actin was visualized with the GFP-tagged Life Actin label. After the addition of Lat A, actin depolymerizes, but centromeres remain clustered and associated with the SPB. (C-E) Quantification of the analysis performed in A and B (see Methods). (C) Scheme of the obtention of centromere (Mis6-GFP) and SPB (Sid4-mCherry) intensity profiles. (D) Calculation of the distance between the centromere and SPB dots. (E) Quantification of the distance between centromeres and SPB in interphase with and without MBC or Lat A. (F) Schematic of the centromere-SPB organization during interphase in *wt* and *sad1.2* cells, at 32°C and 36°C. ONM, outer nuclear membrane; INM, inner nuclear membrane. (G) Quantification of centromere-SPB dissociations. Tags as in Figure 1; centromeres in *wt* and *sad1.2* settings were visualized via Mis6-GFP. Scale bar means 5 μm. N = 50 for each genotype. No asterisk depicts no statistically significant difference. Fisher’s exact test: ****, p < 0.0001. (H) Drop dilution-assays. Sensitivity of the different strains analyzed to chronic treatment with MBC. Serial dilutions (5fold) of normalized exponentially growing cultures were spotted onto YES plates containing DMSO (control) or different amounts of MBC, as indicated, and incubated at 32°C for 48 h.

Similarly, we next wondered about the possible role of actin in maintaining the Rabl configuration, given its role as a motor that promotes the telomere positioning at the NE during budding yeast meiotic prophase (Trelles-Sticken et al. 2005). To explore the role of actin in kinetochore-SPB associations in fission yeast, we disrupted actin with the addition of Latrunculin A (Lat A), an actin polymerization inhibitor, to exponentially growing cells. As control of actin disruption, we used an *S. pombe* strain harboring a GFP-tagged version of Lifeact (LA-GFP), a peptide that labels F-actin *in vivo* (Riedl et al. 2008; Huang et al. 2012). After 10 min of treatment, we observed major actin structural defects visualized by LA-GFP in wild-type-treated cells compared to untreated (Figure 2B). However, under these conditions, we noticed that centromere-SPB associations persist at the same level than in a *wt* untreated setting (Figure 2C–2E). Hence, we discard a major role of actin in the Rabl configuration maintenance, discarding the possibility of using this approach to develop a Rabl configuration-deficient scenario.

To sum up, our results showed that the centromere-SPB interactions, and consequently, the Rabl chromosome configuration, seem to be independent of microtubules differing from the situation found in budding yeast, also independent of actin. This suggests that the Rabl nuclear organization in *Saccharomyces cerevisiae* might be more dynamic than in *Schizosaccharomyces pombe*, which seems to be rather more fixed. In fact, in contrast to fission yeast, it has been observed that efficient DNA damage repair promotion needs centromeres disconnection from the SPB in budding yeast, which depends on microtubule dynamics (Strecker et al. 2016).

### Phosphomutant and phosphomimetic versions of Sad1 residues Thr-3 and Ser-52 do not lead to centromere dissociation defects

An independent approach to address the complete disruption of the Rabl configuration in *S. pombe* is to use the thermo-sensitive Sad1 allele, *sad1.2*, with which all centromeres dissociate from the SPB when *sad1.2* cells are growth at 36°C. However, this scenario leads to total cell lethality due to a failure in the SPB insertion into the NE and spindle formation (Fernandez-Alvarez et al. 2016). Conversely, the growth of this strain at semi-permissive temperature (32°C) produces only partial centromere-SPB defects and does not completely disrupt the Rabl configuration (Fernandez-Alvarez and Cooper 2017b) (Figure 2F). Hence, we tried to generate more penetrant versions of the *sad1.2* allele to promote a greater centromere-SPB disassociation phenotype. The Sad1.2 protein version harbors two single Thr-3-Ser and Ser-52-Pro substitutions, being Ser-52 a validated phosphorylation site for the cyclin-dependent protein kinase Cdc2/CDK-1 (Fernandez-Alvarez et al. 2016; Carpy et al. 2014; Swaffer et al. 2016). We generated phosphomutant and phosphomimetic alleles for Thr-3 and Ser-52 residues in cells harboring Sid4-mCherry and mCherry-Atb2 as SPB and tubulin markers, respectively. We could not find centromere-SPB clustering defects in any of the mutants studied except for the previously characterized Sad1.2 (T3S S52P) (Fernandez-Alvarez et al. 2016)(Figure 2G). This analysis also showed that only the combination of T3S S52P leads to centromere declustering from SPB since all the other combinations produce a *wt*-like phenotype, with almost no defects at all. Congruently, analysis of cellular growth on MBC-containing media showed higher hypersensitivity for *sad1.2* cells compared to the other *sad1* mutant allele combinations (Figure 2H). Previous studies have shown that MBC hypersensitivity in mutants showing some degree of centromere-SPB dissociation is associated with problems in spindle recapture of chromosomes during mitosis; the fact that centromeres were located far from the SPB could complicate their capture by microtubules during mitosis, showing a high sensitivity to MBC-induced microtubule loss (Hou et al. 2012). Our observations argue against the possibility that the association of centromeres to SPB was regulated only by phosphorylation of Sad1 residues Thr-3 and Ser-52. Therefore, the ability to control the centromere association with Sad1 must be controlled by other complementary mechanisms that are probably altered by the Thr-3-Ser and Ser-52-Pro substitutions. Current studies aim to decipher the basis for these centromere-SPB associations.

### Loss of Csi1 in *sad1.2* cells leads to a higher rate of total centromere-SPB dissociations

The foregoing observations indicate that the Rabl chromosome configuration in *S. pombe* seems to be less dynamic than in budding yeast, since microtubule depolymerization has no impact on the centromere-SPB interactions. Since the analysis of the *sad1* alleles suggested the role of alternative proteins, we sought to induce the total transient centromere declustering by eliminating the stabilizer of the centromere-Sad1 associations, Csi1, together with the LEM-domain INM protein Lem2 (Hou et al. 2012; Barrales et al. 2016)(see Figure 2F). With this aim, we constructed strains combining deletions of all the three genes that encoded for Sad1, Csi1, and Lem2 proteins. The double deletion of *csi1* and *lem2* on a *sad1.2* setting leads, in most cases, to spores unable to germinate, indicating a defective cell growth of the triple mutant (Figure 3A). Sporadically, *sad1.2 lem2*Δ *csi1*Δ cells managed to grow, probably associated with the compensatory increased *lnp1* gene expression, as previously observed (Tange et al. 2016). Due to these severe viability defects, we discarded to work with the triple mutant. Analysis of the behavior of all possible double mutant combinations showed that double loss of Csi1 and Lem2 severely impact on cell viability. These defects have been associated with defective pericentromeric heterochromatin identity, which impairs kinetochore proteins association to centromeres, leading to chromosome loss and subsequent growing defects on MBC-containing media, as previously has been reported (Barrales et al. 2016; Hou et al. 2012) (Figure 3B and 3C). The other combination showing a major defect on cellular growth was the double mutant *sad1.2 csi1*Δ. (Figure 3B); these cells also showed increased sensitivity to MBC (Figure 3C). Thus, these experiments pointed out the double mutants *sad1.2 csi1*Δ and *lem2*Δ *csi1*Δ as possible scenarios with greater centromere-SPB dissociation. It has been reported that 10-15% of *lem2*Δ *csi1*Δ cells show all centromeres transitorily disconnected from the SPB (Barrales et al. 2016; Fernandez-Alvarez and Cooper 2017b), but no information has been obtained yet about the double mutant *sad1.2 csi1*Δ. For this reason, we investigated the state of the centromere-SPB contacts in *sad1.2 csi1*Δ cells using live fluorescence microscopy. For a comparative purpose, we included all the strains generated and allocated centromere dissociation phenotypes into two categories: i) *Partial centromere-SPB dissociation*, when, at least, one centromere is detached from the SPB during the analysis (example in −40’ frame in Figure 3E, quantitation in Figure 3G); and ii) *Total centromere-SPB dissociation* when all three centromeres are dissociated from the SPB. In this last category, we established two subtypes: *transient*, at least one frame in interphase during our time-lapse analysis showed total centromere-SPB dissociation (example in −30’ frame in Figure 3E, quantitation in Figure 3H); or *persistent*, similar to the previous one, but centromeres did not interact with the SPB at all at any time during the analysis at interphase (Figure 3F, quantification in Figure 3I). In the case of *transient total centromere dissociation*, cells are still able to divide since one centromere-SPB interaction is sufficient to trigger the SPB insertion into the NE, which allows spindle formation (Fernandez-Alvarez et al. 2016). In contrast, in the *persistent* category, the SPB insertion and, consequently, spindle formation is abolished (Fernandez-Alvarez et al. 2016) (Figure 3F and quantitation in Figure 3I). We assigned the phenotypes of *sad1.2, csi1*Δ, *lem2*Δ, and the double mutant combinations to these categories. Noteworthy, we found more severe defects in *sad1.2 csi1*Δ cells: around 80% of *sad1.2 csi1*Δ cells showed centromere clustering defects, and most importantly, ~25% of this mutant cells showed *transient total centromere dissociation* being this category never seen in the single *csi1*Δ, *lem2*Δ or *sad1.2* single mutants. On the other hand, ~9% of *sad1.2 csi1*Δ cells displayed *persistent total centromere dissociation* reduction in cellular viability (Figures 3B and 3I). Interestingly, although cell growth defects and MBC sensitivity of the *lem2*Δ *csi1*Δ strain are more severe in comparison with the *sad1.2 csi1*Δ genotype (Figures 3B and 3C), the rate and strength of centromere-SPB dissociation of the *lem,2*Δ *csi1*Δ mutant is significantly lower, which suggests that part of the growth defects might be independent of the loss of centromere-SPB contacts; previous works that demonstrated the role of Lem2 in the maintenance of the heterochromatin and nuclear envelope might justify these differences (Kume et al. 2019; Barrales et al. 2016; Tange et al. 2016). In conclusion, we identified *sad1.2 csi1*Δ as an optimal scenario where exploring the behavior of the kinetochore in Rabl chromosome configuration-deficient cells for the combination of two reasons: 1) its higher and severe defects in total centromere-SPB dissociation and 2) its lower impact on cell viability compared to the *sad1.2, csi1*Δ, *andlem2*Δ setting.

**Figure 3.**
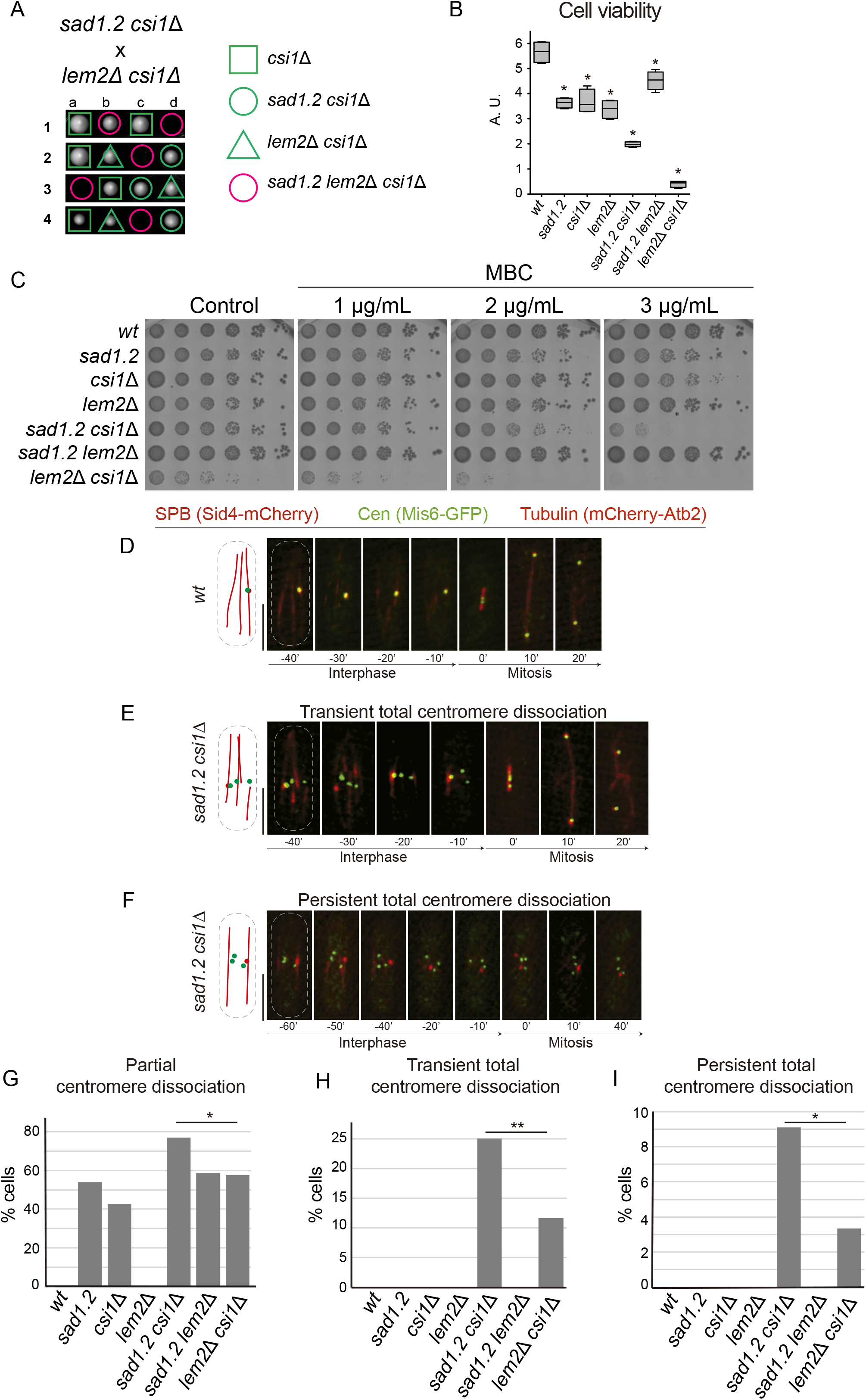
Loss of Csi1 in *sad1.2* cells leads to a higher rate of total centromere dissociation from the SPB without severely compromising cell viability. (A) *sad1.2 lem2*Δ *csi1*Δ cells present a synthetic lethality when spores germinate after tetrads dissection analysis. An example of the *sad1.2 csi1*Δ and *lem2*Δ *csi1*Δ cross is shown. Spores were grown at 32°C for 5 days. (B) Cell viability after combining *csi1* deletion, *lem2* deletion, and/or the presence of the thermosensitive *sad1.2* allele. Cells growing in YES medium for 16 h were diluted in fresh medium during two generation times, normalized to 6 x 10^6^ cells /ml, platted onto YES plates, and incubated for 72 h at 32°C (N = 4). Fisher’s exact test was used to determine *p-values* between wild-type and mutant strains; *p < 0.05. (C) Drop dilution-assays. Sensitivity of the different analyzed strains to chronic treatment with MBC. Serial dilutions (5-fold) of normalized exponentially growing cultures were spotted as indicated in Figure 2H. (D-F) Frames from films of proliferating cells; SPBs and spindles are visualized as in Figure 1. Centromeres were visualized via Mis6-GFP. Scale bar represents 5 μm. (G-I) Quantification of the phenotypes shown in D-F (see main text for details), in each genotype scoring more than 30 cells in, at least, three independent experiments. *p-values* between *sad1.2 csi1*Δ and *lem2 csi1*Δ were determined by Fisher’s exact test; **p < 0.01, *p < 0.05.

### Ndc80 and Nuf2 dislocate from centromeres during interphase in Rabl deficient cells

To further elucidate the behavior of the kinetochore when it is not associated with the SPB during interphase, we analyzed inner and outer kinetochore proteins at the centromeres in cells with and without the Rabl chromosome conformation. We tested Cnp20 and Mis6 as canonical inner kinetochore proteins and Ndc80 and Nuf2 as outer kinetochore proteins. With this aim, we constructed *sad1.2 csi1*Δ strains harboring endogenously GFP-tagged Cnp20, Ndc80, and Nuf2 together with the previously analyzed Mis6. Consistently with our previous observations with Mis6-GFP (Figure 3D–3F), we found that inner kinetochore protein Cnp20 showed normal location at the centromeres during interphase in *wt* (Figure 4A) as well as *sad1.2 csi1*Δ cells, even though when these are dissociated from the SPB (Figure 4B). In contrast, we noticed that the Ndc80-GFP and Nuf2-GFP signals are absent at the centromeres when these are totally dissociated from the SPB during interphase (Figure 4D and Supplementary Figure 1). The outer kinetochore complex is probably not stable at the centromeres in interphase without the interaction with the SPB. This is important because, so far, it was believed that loss of Ndc80 or Nuf2 leads to centromere declustering as naturally occurs in meiosis (Asakawa et al. 2005); however, both proteins require their interaction with the SPB to be persistently associated to the centromeres. We hypothesized that, once Ndc80 and Nuf2 are dislocated from the centromere due to the absence of contact with the SPB, this centromere will not be able to interact more and will be declustered from the SPB until miotic onset. Current studies aim to decipher the basis for the refunding of centromere-SPB interactions after a dissociation. We did not find this phenotype in the *sad1.2* or *csi1*Δ single mutants, meaning that the disassembly of outer kinetochore components requires, at least, *transient* total centromere-SPB dissociation (Figure 4E). Thus, together with the previous observation that the loss of the kinetochore proteins impacts on the Rabl configuration, our results now suggest that this relationship is bidirectional: removing the Rabl configuration by mutation of the NE proteins Csi1 and Sad1 causes the outer kinetochore proteins to disperse from the centromeres in interphase. Moreover, the fact that *sad1.2* and *csi1*Δ single mutants at normal growth conditions (32°C) always show at least one centromere interaction with the SPB is probably enough to maintain the outer kinetochore structure since the interaction between centromeres and SPB is dynamic, and centromeres tend to be permanently associated to the SPB during fission yeast interphase. Hence, if the centromere is permanently dissociated from the SPB for enough time, it is likely that the outer kinetochore losses stability or biochemical signals to continue associated with the centromere, being disassembled. This situation only could occur when all centromeres are declustered from the SPB (transiently or permanently), which occurs in *sad1.2 csi1*Δ cells but not in the *sad1.2 or csi1*Δ single mutants. The role of the centromeric region in the SPB insertion into the NE (Fernandez-Alvarez et al. 2016), a yeast specific mechanism, might be the reason for the conservation of the Rabl configuration in fission yeast and, consequently, the preservation of the outer kinetochore structure throughout the cell cycle.

**Figure 4.**
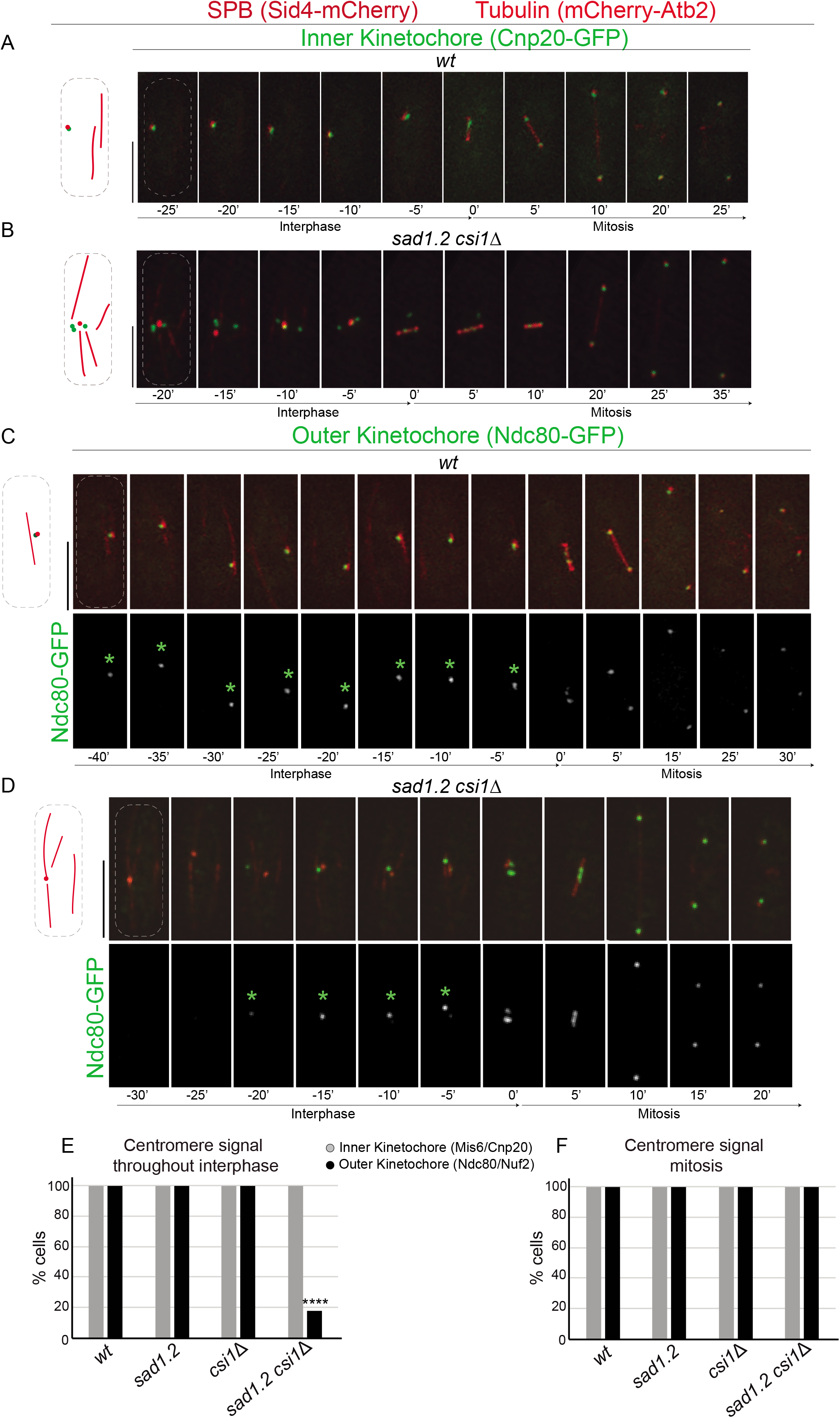
Outer kinetochore component Ndc80 is disassembled in interphase and assembled at mitotic onset in Rabl-deficient cells. (A-D) Frames from films of mitotic cells; tags and numbers as in Figure 1. Scale bars represent 5 μm. Green asterisks indicate the presence of the Ndc80 signal. (E-F) Quantification of the centromere signals (tags as in Figure 1); N=50 for each genotype with more than three independent experiments. *p-value* was determined by Fisher’s exact test, **** p < 0.0001. Cells were scored as negative when the kinetochore signal was missed during all interphase frames and was recovered around 20 min before mitosis.

### The outer kinetochore accumulates at the mitotic onset

During the analysis, we noticed that although Ndc80 and Nuf2 signals are absent in interphase *sad1.2 csi1*Δ cells, these proteins are located at the centromeres in mitotic cells, which is easily recognizable by the presence of two SPBs (Figure 4 and Supplementary Figure 1). This observation indicated that the outer kinetochore was able to be reconstructed at the mitotic onset to prepare the cells for chromosome segregation. In more detail, we found that Ndc80-GFP and Nuf2-GFP signals accumulate during late prophase 20-30 min before SPB separation, gradually increasing until reaching similar levels to *wt* cells. This accumulation of Ndc80 at the centromeres is never seen in a *wt* setting (Figure 1) and precedes the later increment of the protein observed during anaphase (Dhatchinamoorthy et al. 2017). Hence, the outer kinetochore, or at least, Ndc80 and Nuf2, two core proteins of the structure, are actively accumulated at mitotic onset in fission yeast in a similar manner, in terms of the timing, to those seen in metazoan. The reason why this mechanism has not been observed before is that the Rabl configuration masks it. A plausible explanation is that the maintenance of the Rabl configuration during interphase appeared in evolution later to the disassembly and reassembly of the outer kinetochore complex. According to this hypothesis, the Rabl configuration function of controlling the SPB insertion into the NE, a yeast-specific mechanism, favors that Ndc80 and Nuf2 stay stable at the centromeres to maintain centromere-SPB interactions. Using our approach, involving the removal of the Rabl configuration, the outer kinetochore is disassembled, but the program to accumulate these proteins at the centromeres in the G2/M transition is conserved and triggered as in metazoans. In fact, the controlled accumulation of the outer kinetochore proteins preceding chromosome segregation is naturally active in fission yeast meiosis, since Ndc80 and Nuf2 are accumulated around 20-30 min before meiosis I (Asakawa et al. 2005). In meiosis, the Rabl conformation is substituted by the telomere bouquet, a meiotic prophase-specific conformation where the telomeres cluster together at the SPB; during this stage, centromeres are not associated to the SPB, in a similar scenario to that seen in our system using the double mutation *sad1.2 csi1*Δ. We think that fission yeast could reuse this conserved program in mitosis, mimicking the meiotic scenario.

Here we showed evidence of unexpected interphase dispersion and pre-mitotic gradual accumulation of two of the main protein complexes, which are integral elements of the outer kinetochore. So far, it has been known that Ndc80 and Nuf2 protein levels at the centromeres were maintained throughout mitotic interphase. Astonishingly, our data suggest that the mechanisms controlling the disassembly and reassembly of the outer kinetochore might also be conserved in fission yeast. Disclosing the existence of this mechanism in *S. pombe* opens up the possibility of future studies using this yeast model to explore the mammalian kinetochore disassembly/assembly program.

**Figure 5.**
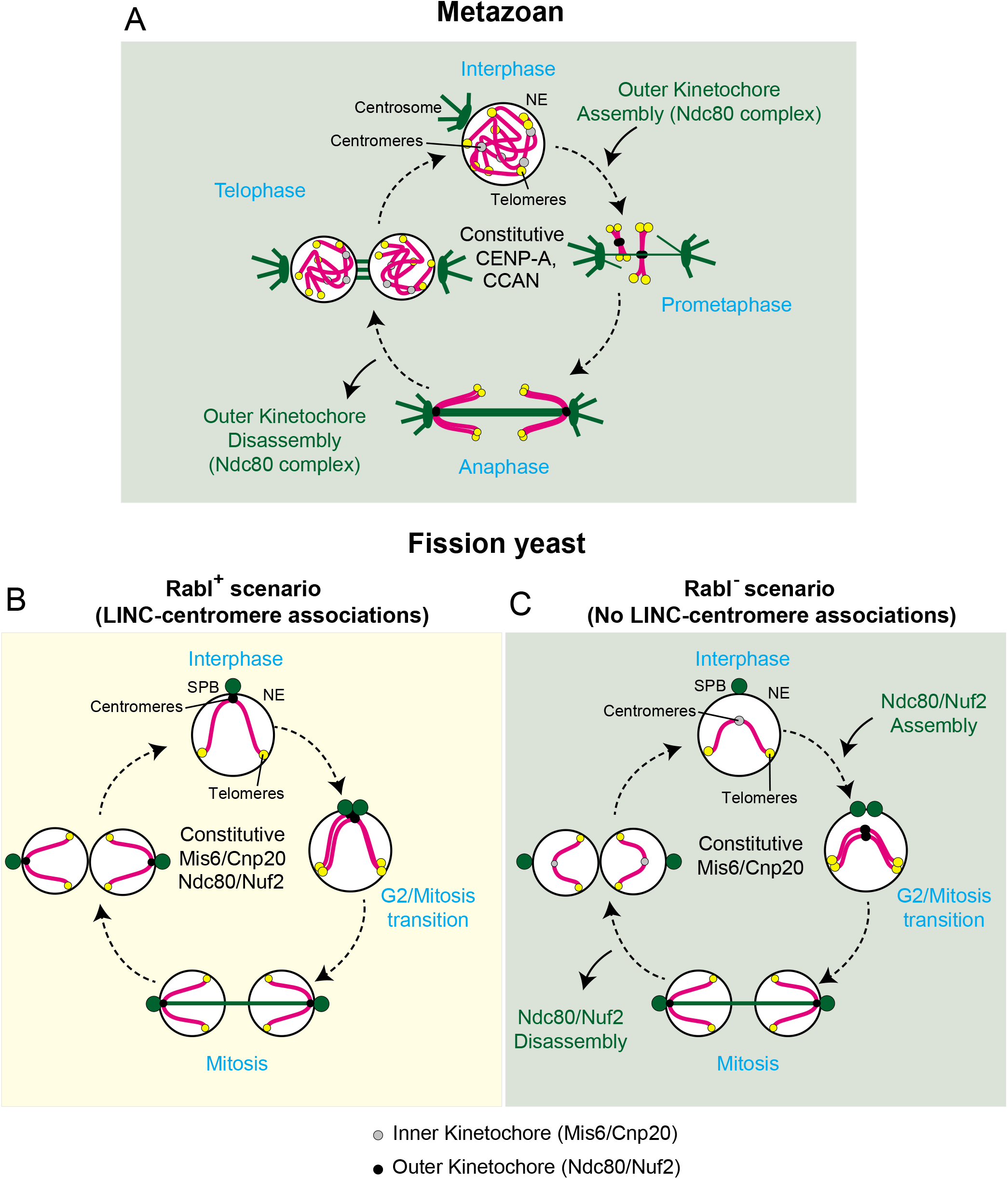
Outer kinetochore delocalization and localization in Rabl-deficient cells during the cell cycle remind the scenario in metazoan. Cell cycle progression in metazoan and fission yeast with and without interphase Rabl chromosome configuration. (A) In metazoan, CENP-A and CCAN (constitutive centromere-associated network) are constitutively associated with centromeres during the cell cycle. Ndc80 complex is assembled and accumulated at centromeres in prophase, and it is delocalized during late anaphase-telophase. (B) In fission yeast, interphase Rabl chromosome configuration requires the constitutive localization of Ndc80 at centromeres. (C) Abolition of the Rabl configuration discloses a delocalization and accumulation cycle of Ndc80 in fission yeast, similar to metazoan.

---

**Supplementary Figure 1. Additional views of the delocalization and accumulation of the outer kinetochore in Rabl deficient cells (Supplement to Figure 4).**

(A-B) Ndc80-GFP and (C-D) Nuf2-GFP signals at the SPB during mitotic interphase are lost with *sad1.2 csi1*Δ mutations. Tags and numbering as in Figure 1. Scale bars represent 5 μm. Green asterisks indicate the presence of Ndc80-GFP or Nuf2-GFP signals.

## METHODS

### Strains and growth conditions

Strains’ growth conditions and molecular biology approaches (Moreno, Klar, and Nurse 1991) were used. Gene deletion and C-terminal tagging were performed as described (Bahler et al. 1998; Fennell et al. 2015). pFA6a plasmids were used to amplify kanMX6, hphMX6, and natMX6 resistance cassettes. Insertions of mCherry-Atb2 at the *aur1* locus (Hashida-Okado et al. 1998) utilized pYC19-mCherryAtb2 (Nakamura et al. 2011) provided by T. Toda (Hiroshima University). Expand Long Template polymerase (Roche) was used for PCR. Haploid cells were usually grown at 32 °C in YE4S or EMM media. Final concentrations of aureobasidin A (0.5 μg/mL), nourseothricin (100 μg/mL clonNAT), G418 (150 μg/mL geneticin) and hygromycin B (300 μg/mL) were added for selection purpose. Strains used in this study are listed in Supplementary Table 1.

#### Sensitivity assays

Strains were revived in solid YES medium, then aerobically precultured up to saturation (D.O. = 1) and subcultured in YES liquid medium with 180 rpm agitation, until D.O. = 0.5 - 0.7 is reached (interphase). Cell viability of normalized exponentially growing cell cultures to 6 x 10^6^ cells/ml was determined by plating cells in triplicate onto YE4S plates and counting colony-forming units after five days incubation at 32°C. For drop-dilution assays, cells growing exponentially at 32°C were normalized to 6 x 10^6^ cells/ml, and 5-fold serial dilutions were spotted onto YE4S plates containing different concentrations of MBC. The plates were incubated at 32°C for 48-72 h.

### Fluorescence microscopy, live analysis, and quantification

Fluorescence microscopy images were generated using the DeltaVision microscope system (Applied Precision, Seattle, WA). Cells were adhered to 35 mm glass culture dishes (MatTek) using 0.2 mg/ml soybean lectin (Sigma) and immersed in EMM (with required supplements). Time-lapse imaging was carried out at 27 °C in an Environmental Chamber with a DeltaVision Spectris (Applied Precision) comprising an Olympus IX70 widefield inverted epifluorescence microscope, an Olympus UPlanSapo 100x NA 1.4 oil immersion objective, and a Photometrics CCD CoolSnap HQ camera. Images were acquired over 26 focal planes at a 0.35 μm step size. For the quantification of protein fluorescence intensity, maximum-projected raw microscopy data were corrected for photo-bleaching via the Exponential Fitting method. Foci intensity time-series were obtained after detection with a Laplacian of Gaussian filter and tracking with the LAP algorithm (TrackMate). Tracks were time-aligned according to splitting events, and intensities were normalized respect to background mean intensity. Images were further deconvolved and combined into a 2D image using the maximum intensity projection setting using softWoRx (Applied Precision). Image processing and analysis were performed using Adobe Photoshop 2020.

### Carbendazim and latrunculin treatments

For carbendazim treatment, a working solution of YES+MBC (15 μg/mL) (carbendazim, CAS No. 10605-21-7) is prepared using a stock solution of DMSO+MBC (2.5 mg/mL). Strains were revived in solid YES medium, then aerobically precultured up to saturation (D.O. = 1) and subcultured, in both cases in YES liquid medium with 180 rpm agitation, until D.O. = 0.3 - 0.4 is reached (interphase). 50 μL of lectin (0.2 μg/mL) (Sigma Aldrich, L1395) is used for cell immobilization on a μ-Slide 8 Well Uncoated (ibidi GmbH). YES+MBC (experiment) or YES (control) medium is used for filming cells.

For latrunculin A treatment, exponentially growing cells were incubated 10 min in 3 mL of YES rich medium with a total concentration of latrunculin A of 6 μM (15 μL of a 1 mM stock). After incubation, cells are immobilized with lectin as in MBC treatment on a coverslip and mounted into a microscope slide (Linealab) for image acquisition.

For live microscopy, images were taken with 100 ms and 50 ms of exposure time for fluorescent and brightfield channels, respectively, and 13 focal planes with a 0.5 μm step size, using a spinning disk confocal microscopy system (Photometrics Evolve camera; Olympus 100x 1.4 NA oil immersion objective; Roper Scientific). For the co-localization analysis, maximum Z-projection images of interphase cells, those with one single Sid4-mCherry dot (SPB), were subjected to co-localization analysis. For each cell, an axis containing the center of both Sid4-mCherry and Mis6-GFP (centromeres) dots is drawn, and the intensity of the pixels from both channels is measured, normalized and plotted along such axis. The resultant intensity profiles are used to measure the distance between the dots, defined as the distance in microns between the x-coordinates of the intensity maxima of both profiles, considered to correspond with the center of the dots.

## Acknowledgments

We thank all lab members for critical comments on the manuscript; Alejandra Cano for technical support; and the CABD microscopy facility technician Katherina García. We would like to thank the Genetics Department and Springboard lab for their useful discussion and comments, especially Víctor Carranco for technical support. This work was supported by the Spanish Government, Plan Nacional project PGC2018-098118-A-I00 and Ramon y Cajal program, RyC-2016-19659 to AF-A; by the Pablo de Olavide University “Ayuda Puente Predoctoral” fellowship (PPI1803) to AP-S; and by the Spanish Education and Professional Formation Ministry, Research Collaboration Grant to DL-P. The CABD is an institution funded by Pablo de Olavide University, Consejo Superior de Investigaciones Científicas (CSIC), and Junta de Andalucía.

## Author contributions

AF-A. designed the study; AJ-M and AP-S. performed most of the experiments with the support of DL-P. DD-G and LM-T contributed to Figure 2; AF-A. acquired funding and supervised the project; AF-A. wrote the manuscript with support of AJ-M, AP-S, and DL-P.

## References

Asakawa, H., A. Hayashi, T. Haraguchi, and Y. Hiraoka. 2005. ‘Dissociation of the Nuf2-Ndc8o complex releases centromeres from the spindle-pole body during meiotic prophase in fission yeast’, Mol Biol Cell, 16: 2325–38.

Bahler, J., J. Q. Wu, M. S. Longtine, N. G. Shah, A. McKenzie, 3rd, A. B. Steever, A. Wach, P. Philippsen, and J. R. Pringle. 1998. ‘Heterologous modules for efficient and versatile PCR-based gene targeting in Schizosaccharomyces pombe’, Yeast, 14: 943–51.

Barrales, R. R., M. Forn, P. R. Georgescu, Z. Sarkadi, and S. Braun. 2016. ‘Control of heterochromatin localization and silencing by the nuclear membrane protein Lem2’, Genes Dev, 30:133–48.

Biggins, S. 2013. ‘The composition, functions, and regulation of the budding yeast kinetochore’, Genetics, 194: 817–46.

Bystricky, K., P. Heun, L. Gehlen, J. Langowski, and S. M. Gasser. 2004. ‘Long-range compaction and flexibility of interphase chromatin in budding yeast analyzed by high-resolution imaging techniques’, Proc Natl Acad Sci U S A, 101:16495–500.

Carpy, A., K. Krug, S. Graf, A. Koch, S. Popic, S. Hauf, and B. Macek. 2014. ‘Absolute proteome and phosphoproteome dynamics during the cell cycle of Schizosaccharomyces pombe (Fission Yeast)’, Mol Cell Proteomics, 13:1925–36.

Cheeseman, I. M. 2014. ‘The kinetochore’, Cold Spring Harb Perspect Biol, 6: aoi5826.

Cheeseman, I. M., C. Brew, M. Wolyniak, A. Desai, S. Anderson, N. Muster, J. R. Yates, T. C. Huffaker, D. G. Drubin, and G. Barnes. 2001. ‘Implication of a novel multiprotein Damip complex in outer kinetochore function’, J Cell Biol, 155:1137–45.

Cheeseman, I. M., and A. Desai. 2008. ‘Molecular architecture of the kinetochore-microtubule interface’, Nat Rev Mol Cell Biol, 9:33–46.

Czapiewski, R., M. I. Robson, and E. C. Schirmer. 2016. ‘Anchoring a Leviathan: How the Nuclear Membrane Tethers the Genome’, Front Genet, ~ŗ. 82.

Dhatchinamoorthy, K., M. Mattingly, and J. L. Gerton. 2018. ‘Regulation of kinetochore configuration during mitosis’, Curr Genet, 64:1197–203.

Dhatchinamoorthy, K., M. Shivaraju, J. J. Lange, B. Rubinstein, J. R. Unruh, B. D. Slaughter, and J. L. Gerton. 2017. ‘Structural plasticity of the living kinetochore’, J Cell Biol, 216: 355i–7¤o

Fennell, A., A. Fernandez-Alvarez, K. Tomita, and J. P. Cooper. 2015. ‘Telomeres and centromeres have interchangeable roles in promoting meiotic spindle formation’, J Cell Biol, 208: 415–28.

Fernandez-Alvarez, A., C. Bez, E. T. O’Toole, M. Morphew, and J. P. Cooper. 2016. ‘Mitotic Nuclear Envelope Breakdown and Spindle Nucleation Are Controlled by Interphase Contacts between Centromeres and the Nuclear Envelope’, Dev Cell, 39: 544–59.

Fernandez-Alvarez, A., and J. P. Cooper. 2017a. ‘Chromosomes Orchestrate Their Own Liberation: Nuclear Envelope Disassembly’, Trends Cell Biol, 27: 255–65.

Fernandez-Alvarez, A., and J. P. Cooper. 2017b. ‘The functionally elusive Rabl chromosome configuration directly regulates nuclear membrane remodeling at mitotic onset’, Cell Cycle, 16:1392–96.

Funabiki, H., I. Hagan, S. Uzawa, and M. Yanagida. 1993. ‘Cell cycle-dependent specific positioning and clustering of centromeres and telomeres in fission yeast’, J Cell Biol, 121: 961–76.

Gu, Y., C. Yam, and S. Oliferenko. 2012. ‘Divergence of mitotic strategies in fission yeasts’, Nucleus, 3: 220–5.

Hagan, I., and M. Yanagida. 1995. ‘The product of the spindle formation gene sadi÷ associates with the fission yeast spindle pole body and is essential for viability’, J Cell Biol, 129:1033–47.

Hara, M., and T. Fukagawa. 2018. ‘Kinetochore assembly and disassembly during mitotic entry and exit’, Curr Opin Cell Biol, 52: 73–81.

Hashida-Okado, T., R. Yasumoto, M. Endo, K. Takesako, and I. Kato. 1998. ‘Isolation and characterization of the aureobasidin A-resistant gene, aur1R, on Schizosaccharomyces pombe: roles of Aur1p+ in cell morphogenesis’, Curr Genet, 33:38–45.

Hattersley, N., D. Cheerambathur, M. Moyle, M. Stefanutti, A. Richardson, K. Y. Lee, J. Dumont, K. Oegema, and A. Desai. 2016. ‘A Nucleoporin Docks Protein Phosphatase 1 to Direct Meiotic Chromosome Segregation and Nuclear Assembly’, Dev Cell, 38: 463–77.

Hiraoka, Y., and A. F. Dernburg. 2009. ‘The SUN rises on meiotic chromosome dynamics’, Dev Cell, τj-. 598–605.

Hou, H., Z. Zhou, Y. Wang, J. Wang, S. P. Kallgren, T. Kurchuk, E. A. Miller, F. Chang, and S. Jia. 2012. ‘Csi1 links centromeres to the nuclear envelope for centromere clustering’, J Cell Biol, 199:735–44.

Hsu, K. S., and T. Toda. 2011. ‘Ndc80 internal loop interacts with Dis1/TOG to ensure proper kinetochore-spindle attachment in fission yeast’, CurrBiol, 21: 214–20.

Huang, J., Y. Huang, H. Yu, D. Subramanian, A. Padmanabhan, R. Thadani, Y. Tao, X. Tang, R. Wedlich-Soldner, and M. K. Balasubramanian. 2012. ‘Nonmedially assembled F-actin cables incorporate into the actomyosin ring in fission yeast’, J Cell Biol, 199: 831–47.

Janke, C., J. Ortiz, T. U. Tanaka, J. Lechner, and E. Schiebel. 2002. ‘Four new subunits of the Dami-Duoi complex reveal novel functions in sister kinetochore biorientation’, EMBO J, 21:181–93.

Jin, Q., E. Trelles-Sticken, H. Scherthan, and J. Loidl. 1998. ‘Yeast nuclei display prominent centromere clustering that is reduced in nondividing cells and in meiotic prophase’, J Cell Biol, 141: 21–9.

Jin, Q. W., J. Fuchs, and J. Loidl. 2000. ‘Centromere clustering is a major determinant of yeast interphase nuclear organization’, J Cell Sci, 113 (Pt 11): 1903–12.

Kume, K., H. Cantwell, A. Burrell, and P. Nurse. 2019. ‘Nuclear membrane protein Lem2 regulates nuclear size through membrane flow’, Nat Commνn, 10:1871.

Liu, X., I. McLeod, S. Anderson, J. R. Yates, 3rd, and X. He. 2005. ‘Molecular analysis of kinetochore architecture in fission yeast’, EMBO J, 24: 2919–30.

Moreno, S., A. Klar, and P. Nurse. 1991. ‘Molecular genetic analysis of fission yeast Schizosaccharomyces pombe’, Methods Enzymol, 194:795–823.

Nabetani, A., T. Koujin, C. Tsutsumi, T. Haraguchi, and Y. Hiraoka. 2001. ‘Aconserved protein, Nuf2, is implicated in connecting the centromere to the spindle during chromosome segregation: a link between the kinetochore function and the spindle checkpoint’, Chromosoma, 110: 322–34.

Nagpal, H., and T. Fukagawa. 2016. ‘Kinetochore assembly and function through the cell cycle’, Chromosoma, 125: 645–59.

Nakamura, Y., A. Arai, Y. Takebe, and M. Masuda. 2011. ‘A chemical compound for controlled expression of nmt1-driven gene in the fission yeast Schizosaccharomyces pombe’, Anal Biochem, 412:159–64.

Obuse, C., O. Iwasaki, T. Kiyomitsu, G. Goshima, Y. Toyoda, and M. Yanagida. 2004. ‘A conserved Mis12 centromere complex is linked to heterochromatic HPi and outer kinetochore protein Zwint-1’, Nat Cell Biol, 6:1135–41.

Riedl, J., A. H. Crevenna, K. Kessenbrock, J. H. Yu, D. Neukirchen, M. Bista, F. Bradke, D. Jenne, T. A. Holak, Z. Werb, M. Sixt, and R. Wedlich-Soldner. 2008. ‘Lifeact: a versatile marker to visualize F-actin’, Nat Methods, 5:605–7.

Shimanuki, M., F. Miki, D. Q. Ding, Y. Chikashige, Y. Hiraoka, T. Horio, and O. Niwa. 1997. ‘A novel fission yeast gene, kmsi÷, is required for the formation of meiotic prophase specific nuclear architecture’, Mol Gen Genet, 254: 238–49.

Strecker, J., G. D. Gupta, W. Zhang, M. Bashkurov, M. C. Landry, L. Pelletier, and D. Durocher. 2016. ‘DNAdamage signalling targets the kinetochore to promote chromatin mobility’, Nat Cell Biol, 18: 281–90.

Swaffer, M. P., A. W. Jones, H. R. Flynn, A. P. Snijders, and P. Nurse. 2016. ‘CDK Substrate Phosphorylation and Ordering the Cell Cycle’, Cell, 167:1750–61 ei6.

Taddei, A., and S. M. Gasser. 2012. ‘Structure and function in the budding yeast nucleus’, Genetics, 192:107–29.

Takahashi, K., E. S. Chen, and M. Yanagida. 2000. ‘Requirement of Mis6 centromere connector for localizing a CENP-A-like protein in fission yeast’, Science, 288: 2215–9.

Tange, Y., Y. Chikashige, S. Takahata, K. Kawakami, M. Higashi, C. Mori, T. Kojidani, Y. Hirano, H. Asakawa, Y. Murakami, T. Haraguchi, and Y. Hiraoka. 2016. ‘Inner nuclear membrane protein Lem2 augments heterochromatin formation in response to nutritional conditions’, Genes Cells, 21: 812–32.

Therizols, P., T. Duong, B. Dujon, C. Zimmer, and E. Fabre. 2010. ‘Chromosome arm length and nuclear constraints determine the dynamic relationship of yeast subtelomeres’, Proc Natl Acad Sci U S A, 107: 2025–30.

Trelles-Sticken, E., C. Adelfalk, J. Loidl, and H. Scherthan. 2005. ‘Meiotic telomere clustering requires actin for its formation and cohesin for its resolution’, J Cell Biol, 170. 213–23.

Unruh, J. R., B. D. Slaughter, and S. L. Jaspersen. 2018. ‘Functional Analysis of the Yeast LINC Complex Using Fluctuation Spectroscopy and Super-Resolution Imaging’, Methods Mol Biol, 1840:137–61.

Wigge, P. A., and J. V. Kilmartin. 2001. ‘The Ndc80p complex from Saccharomyces cerevisiae contains conserved centromere components and has a function in chromosome segregation’, J Cell Biol, 152: 349–60.

