## Supplementary figures and images for "Metazoan-like kinetochore arrangement masked by the interphase RabI configuration"

### Supplementary Figure 1

Supplementary Figure 1

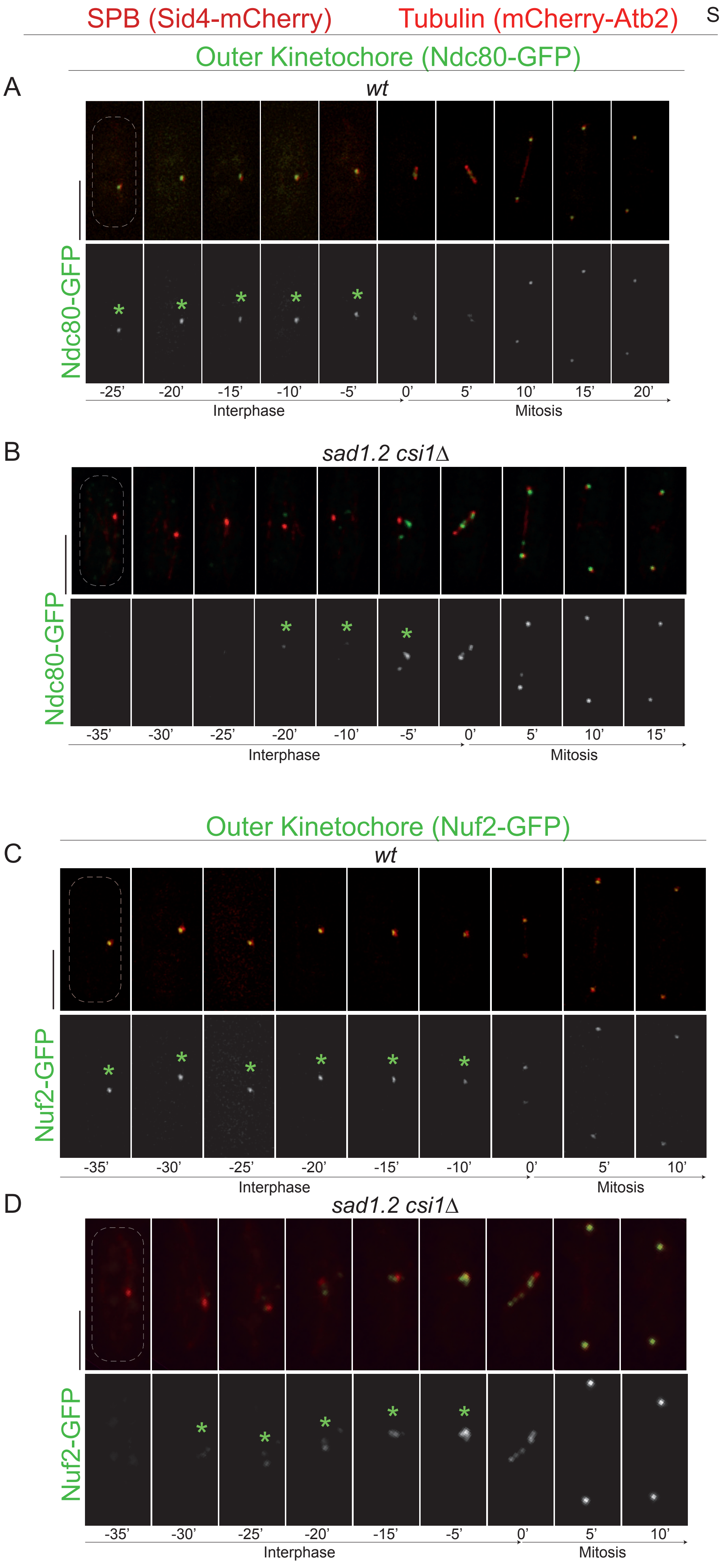
